# A new schizophrenia model: immune activation is associated with induction of the tryptophan catabolite pathway and increased eotaxin levels which together determine memory impairments and schizophrenia symptom dimensions

**DOI:** 10.1101/393173

**Authors:** Sunee Sirivichayakul, Buranee Kanchanatawan, Supaksorn Thika, André F. Carvalho, Michael Maes

## Abstract

**Abstract:** *Objective:* Recently, we reported that stable-phase schizophrenia is characterized by two interrelated symptom dimensions: PHEMN (psychotic, hostility, excitation, mannerism and negative symptoms); and DAPS (depressive, anxiety and physio-somatic symptoms) and that Major Neuro-Cognitive psychosis (MNP) is the full blown phenotype of schizophrenia (largely overlapping with deficit schizophrenia). Herein we examined the effects of immune activation in association with tryptophan catabolite (TRYCAT) patterning and memory disorders on PHEMN/DAPS dimensions and MNP.

*Method:* Serum levels of macrophage inflammatory protein-1 (MIP-1), soluble interleukin (IL)-1 receptor antagonist (sIL-1RA), IL-10, eotaxin, IgA/IgM responses to TRYCATs, and Consortium to Establish a Registry for Alzheimer’s disease (CERAD) tests were assessed in 40 controls and 80 schizophrenia patients.

*Results:* Schizophrenia and MNP were predicted by significantly increased levels of IL-10, eotaxin and TRYCATs. A large part of the variance in both PHEMN/DAPS symptom dimensions (42.8%) was explained by cytokine levels and TRYCATs combined. The MTP+sTL-1R A+IL-10 composite score and eotaxin explained each around 19% of the variance in symptom dimensions, and approximately 18% of memory deficits. Moreover, MIP+sIL-1RA+IL-10 was significantly associated with elevations in picolinic acid, xanthurenic acid and 3-OH-kynurenine. Partial Least Squares path modeling shows that the highly significant effects of MIP+sIL-1RA+IL-10 on symptomatology are mediated by the effects of noxious TRYCATs on memory deficits.

*Conclusions:* Current findings indicate that in schizophrenia, immune activation may underpin activation of indoleamine-2,3-dioxygenase and kynurenine monooxygenase, while impairments in episodic and semantic memory may be caused by the neurotoxic effects of TRYCATs and eotaxin. The combined effects of immune activation, eotaxin and memory defects determine to a large extent PHEMN/DAPS symptoms and the MNP phenotype. These findings indicate that schizophrenia phenomenology is largely mediated by multiple neuro-immune pathways and that immune activation, increased production of eotaxin and neurotoxic TRYCATs (picolinic acid, xanthurenic acid and 3-HO-kynurenine) are new drug targets in schizophrenia and MNP.

## 1. Introduction

Recently, we reported that stable-phase schizophrenia is characterized by two interrelated symptom dimensions, namely a) the PHEMN dimension comprising psychotic, hostility, excitation, mannerism and negative symptoms; and b) the DAPS dimension, comprising depressive, anxiety and physio-somatic symptoms [1]. These findings show that the two-syndrome framework differentiating “negative from positive symptoms” [2] is not accurate because PHEM (psychosis, hostility, excitation, mannerism) and negative symptoms belong to one and the same dimension [1]. Moreover, previous theories suggesting that affective symptoms are differently associated with positive versus negative symptoms [3,4] should be revised because all symptoms of both dimensions load highly on an overarching “generalized psychopathology dimension”, which comprises two inter-related PHEMN and DAPS subdimensions. In addition, negative, affective and physio-somatic symptoms of schizophrenia are strongly predicted by impairments in semantic and episodic memory as measured with Consortium to Establish a Registry for Alzheimer’s disease (CERAD) tests [1,5,6].

Moreover, we found that stable phase schizophrenia may be accurately divided into two distinct nosological entities which are qualitatively different from each other and from control subjects, namely Major Neuro-Cognitive Psychosis (MNP) and Simple Neuro-Cognitive Psychosis (SNP) [7]. Both schizophrenia subtypes are characterized by increased PHEMN and DAPS symptoms and severe impairments in semantic memory, whilst MNP is differentiated from SNP by increased PHEMN and DAPS symptoms, impairments in episodic memory and specific biomarkers [7]. There is a strong association between MNP and the deficit phenotype as defined with SDS criteria [8], although the MNP criteria are more restrictive. Other characteristics of MNP versus SNP are a lowered body mass index (reflecting a leptosome body type of this type of schizophrenia), lowered education (suggesting lower cognitive reserve as a risk factor for MNP) and lowered quality of life [7]. Therefore, MNP is the core illness or full blown phenotype, while SNP is the less exuberant phenotype [7]. Previous classifications of schizophrenia could not be validated using unsupervised learning techniques, including “type 1” (positive) versus “type 2” (negative) schizophrenia [2].

Importantly, PHEMN and DAPS dimensions and the MNP and SNP nosological entities are strongly predicted by neuro-immune changes as indicated by changes in tryptophan catabolite (TRYCAT) pathway patterning [1,7]. Elevated levels of IgA-mediated responses directed against noxious TRYCATs including picolinic acid (PA), xanthurenic acid (XA), 3-OH-kynurenine (3HK) and quinolinic acid (QA) are associated with both dimensions, memory deficits, the diagnosis schizophrenia and especially with MNP [1,7,9]. These findings indicate that the TRYCAT pathway is activated in schizophrenia and MNP and that the neurotoxic and excitotoxic effects of the noxious TRYCATs may at least in part underpin cognitive deficits and different symptomatic dimensions of this illness [1,7,9]. Moreover, impairments in IgM-mediated responses to noxious TRYCATs and the KA / 3HK ratio, indicating changes in immune regulation, were strongly associated with cognitive impairments and additionally separated MNP from SNP as different nosological entities [10].

The TRYCAT pathway may be stimulated during an immune response through activation of indeoleamine-2,3-dioxygenase (IDO) by Thelper (Th) 1 and macrophagic M1 cytokines, including interferon (IFN)-γ, interleukin (IL)-1β and IL-6 [11]. There is now evidence that acute episodes of schizophrenia, chronic schizophrenia and first episode psychoses are accompanied by multiple signs of M1, Th-1, Th-2, and T regulatory (Treg) activation, mild chronic inflammation, a chronic “acute phase” response and increased complement factors [12-17]. Therefore, it may be hypothesized that TRYCAT pathway activation in schizophrenia is the consequence of immune activation. Moreover, some products of Ml and Th-1 cells, including IFN-γ, IL-1β and IL-6, activate IDO and exert neurotoxic effects thereby inducing neurocognitive deficits in animal models [18]. Nevertheless, there are no data in schizophrenia that assessed whether the effects of neurotoxic TRYCATs on schizophrenia phenomenology (including symptoms and cognitive deficits) could be initiated by immune mechanisms and whether immune activation is associated with memory impairments thereby explaining PHEMN and DAPS symptoms and MNP as well.

Hence, the aim of the present study was to examine the effects of M1, i.e. macrophage inflammatory protein (MIP)-1 and soluble IL-1 receptor antagonist (sIL-1RA), Th-2 (IL-10) and eotaxin (CCL11) on TRYCAT pathway activation, cognitive deficits and schizophrenia phenomenology (PHEMN and DAPS, SNP and MNP) and whether the immune effects on schizophrenia phenomenology are mediated by changes in TRYCAT pathway patterning leading to cognitive impairments.

## 2. Methods

### 2.1. Participants

In this case control study we recruited schizophrenia patients and healthy controls from the same catchment area (Bangkok, Thailand), who were all Thai nationals, aged 18 to 65 years and of both sexes. We enrolled 80 consecutively admitted outpatients with schizophrenia fulfilling the diagnostic criteria for schizophrenia of the DSM-IV-TR and who were all in a stable phase of the illness. Patients were admitted to the outpatient clinic of the Department of Psychiatry, Faculty of Medicine, Chulalongkorn University, Bangkok, Thailand. Exclusion criteria for patients were: axis-1 DSM-IV-TR disorders other than schizophrenia, including major depression, bipolar disorder, schizoaffective disorder and substance use disorders; neuroinflammatory disorders, including stroke, Parkinson’s disease and multiple sclerosis; acute psychotic episodes; major medical illness including diabetes (type 1 and 2), rheumatoid arthritis, psoriasis, inflammatory bowel disease and chronic obstructive pulmonary disease. Participants (controls or patients) were excluded when using immunomodulatory drugs and supplements with ω3-polyunsaturated fatty acids and antioxidants. Healthy controls were excluded when they presented with any lifetime diagnosis of DSM-IV-TR axis I mental disorders or with a family history of psychotic disorders.

All participants as well as the guardians of patients (parents or other close family members), provided written informed consent prior to participation in this study. The study was conducted according to Thai and International ethics and privacy laws. Approval for the study was obtained from the Institutional Review Board of the Faculty of Medicine, Chulalongkorn University, Bangkok, Thailand, which is in compliance with the International Guidelines for Human Research protection as required by the Declaration of Helsinki, The Belmont Report, CIOMS Guideline and International Conference on Harmonization in Good Clinical Practice (ICH-GCP).

### 2.2. Clinical assessments

In this study, we assessed socio-demographic, clinical, cognitive, psychopathological and biomarker data. The DSM-IV-TR diagnosis of schizophrenia was made using the Mini-International Neuropsychiatric Interview (M.I.N.I.) in a validated Thai translation [19]. We divided the patients into two groups, namely MNP and SNP using results of unsupervised machine learning as explained in Kanchanatawan et al. [7]. A senior psychiatrist also performed a semistructured interview thereby collecting data on socio-demographics, medical history, family history of psychosis. The same senior psychiatrist also scored a number of rating scales, namely the SDS [8] and SANS [20] to assess negative symptoms; the Positive and Negative Syndrome Scale (PANSS) [21] to measure negative (PANSSneg) and positive (PANSSpos) symptoms; the Brief Psychiatric Rating Scale [22] to assess general psychopathology; the Hamilton Depression (HAMD) and Anxiety (HAMA) Rating Scale [23,24] to measure severity of depressiona and anxiety, respectively; and the 12 item Fibromyalgia and Chronic Fatigue Syndrome Rating scale (FF) [25] to measure physio-somatic symptoms. As explained by Kachanatawan et al. [1] we computed 4 PHEMN dimensions using the BPRS and PANSS to measure z-unit weighted composite scores, namely psychotic symptoms: z score PANSS P1 (delusion) (zP1) + zP3 (hallucinations) + zP6 (suspiciousness) + zBPRS11 (suspiciousness) + zBPRS12 (hallucinatory behavior) + BPRS15 (unusual thought content); hostility: zP7 (hostility) + zPANSS general14 (zG14, poor impulse control) + zBPRS10 (hostility) + zBPRS14 (uncooperativeness); excitement: zP14 (excitement) + zP5 (grandiosity) + zBPRS8 (grandiosity) + zBPRS17 (excitement); mannerism: zG5 + zBPRS7 (both mannerism and posturing). The diagnosis of Tobacco Use Disorder (TUD) was made using DSM-IV-TR criteria. We also measured body mass index (BMI) as body weight (kg) / length (m^2^).

The same day that the rating scales were scored and the semistructured interview was completed we also performed CERAD-Neuropsychological tests [5]. The CERAD was assessed by a well-trained research assistant (ST) who is master in mental health. This research assistant was blinded to the clinical diagnoses and performed all CERAD tests in patients and controls. The CERAD-Neuropsychology battery comprises cognitive tests, which assess several domains including episodic and semantic memory and general neuropsychological functioning. Included are: <u>Mini-Mental State Examination</u> (MMSE), to assess different functions such as concentration, orientation, naming, constructional praxis and memory. <u>Verbal Fluency Test</u> (VFT), to assess semantic memory, language, fluency, cognitive flexibility and verbal productivity. <u>Word List Memory</u> (WLM) to measure verbal episodic memory, learning ability and working memory for verbal information. <u>Word List Recall, true recall</u> (WLR True) to probe verbal episodic memory–recall. <u>Word List Recall, false recall</u> (WLR False) to assess intrusion errors or false memory creation. In addition, we used two indices of episodic and semantic memory based on two oblimin-rotated principal components (PCs) extracted from 8 CERAD tests and explaining 68.3% of the variance: a) the first semantic memory PC loading highly on VFT, Boston naming Tests, MMSE and Constructive Praxis (reflecting impairments in semantic memory, naming and a generalized cognitive impairment); and b) the second episodic memory PC loading highly on WLM, WLR True, WLR False and Word List Recognition [6].

### 2.3. Biomarker assays

The same day that we performed the interviews and completed CERAD tests, fasting blood was sampled around 8.00 a.m. and frozen at −80 ^o^C until thawed for assay of biomarkers. For cytokines/chemokines, 50 μl of serum (1:2 dilution in calibrator diluent) was mixed with 50 μl of microparticle cocktail containing eotaxin, sIL-1ra, IL-10 and MIP-1α (R&D Systems, Inc, Minneapolis, MN, USA) per well of a 96-well plate provided by manufacturer and incubated for 2 hours at room temperature on a shaker at 800 rpm. The mixture was then washed 3 times with wash buffer and 50 μl diluted Biotin Antibody cocktail was added and then incubated for 1 hour. Wells were washed 3 times before another 50 μl of diluted Streptavidin-PE was added and further incubated for 30 minutes. Finally, wells were washed 3 times and 100 μl of wash buffer was added and left at room temperature for 2 minutes before being read with Bio-Plex^®^ 200 System (Bio-Rad Laboratories, Inc.). The intra-assay CV values were <7.0%. The least detectable dose was 1.82 pg/mL for eotaxin, 1.58 pg/mL for MIP-1, 5.98 pg/mL for IL-1RA and 0.4 pg/mL for IL-10. “The 6 TRYCATs were assayed as described before [26,27]. Briefly, TRYCATs were dissolved in 200 μL dimethylsulfoxide (DMSO) (Acros). Bovine serum albumin (BSA) (ID Bio) was dissolved in 3mL 2-morpholino-ethanesulfonic acid monohydrate (MES Acros) buffer 10–1 M at pH = 6.3 (Acros). The TRYCATs were then mixed with the BSA solution and supplemented with 15 mg N-hydroxysuccinimide (Sigma) and 1-(3-dimethylaminopropyl)-3-ethylcarbodiimide (Acros) as coupling agents. The conjugates were synthesized by linking 3HK (Sigma), KA (Acros), QA (Acros), AA (Acros), XA (Acros) and PA (Acros) to 20 mg BSA. The coupling reaction proceeded at 37°C for 1 hour in the dark. The coupling was stopped by adding 100 mg hydroxylamine (Sigma-Aldrich) per conjugate. Protein conjugates were dialyzed with 10–1 M NaCl solution, pH=6 for 72 hours, with the bath solution being changed at least four times per day. The conjugated TRYCATs and BSA concentrations were evaluated by spectrophotometry. The coupling ratio of each conjugate was determined by measuring the concentration of TRYCATs and BSA at 310–330 nm and 280 nm, respectively. ELISA tests were used to determine plasma titers of serum immunoglobulin (Ig)M and IgA. Towards this end, polystyrene 96-well plates (NUNC) were coated with 200 μL solution containing 10-50 μg/mL TRYCAT conjugates in 0.05 M carbonate buffer (pH = 9.6). Well plates were incubated under agitation at 4 °C for 16 hours. Then, 200 μL blocking buffer A (Phosphate Buffered Saline, PBS, 2.5 g/L BSA, pH=7) was applied and all samples were incubated at 37 °C for 1 hour. Well plates were washed with PBS solution and filled with 100 μL serum diluted 1:130 in blocking buffer and incubated at 37 °C for 1 hour and 45 minutes. Well plates were washed 3 times with PBS, 0.05% Tween 20, incubated with peroxidase-labeled goat anti-human IgA (SouthernBiotech) antibodies at 37 °C for 1 hour. The goat anti-human IgM antibody was diluted at 1:5000 and the IgA antibody was diluted at 1:10,000 in blocking buffer (PBS, 2.5 g/L BSA). Plates were then washed three times with PBS, 0.05% Tween 20. Fifty μL of 3,3′ ,5,5′-Tetramethylbenzidine (TMB) substrate (SouthernBiotech) was added and incubated for 10 minutes in the dark. The reaction was stopped using 50 μl of TMB stop solution (SouthernBiotech). Optical densities (ODs) were measured at 450 nm using Varioskan Flash (Thermo Scientific). All assays were carried out in duplicate. The analytical intra-assays CV values were < 7%. The OD scores were expressed as z scores”. We computed 3 z-unit weighted composite scores as explained previously (Kanchanatawan et al., 2018b; 2018c): a) zIgA NOX_PRO = sum of z scores of QA (zQA) + zPA + zXA – zAA – zKA (index of increased noxious potential); b) ΔNOX_PRO = zIgA (zQA + zPA + zXA +z3HK – zAA – zKA) – zIgM (zQA + zPA + zXA +3HK – zAA – zKA) (a more comprehensive index of increased noxious potential); and c_ zIgM KA_3HK = zIgM KA – z3HK (index of lowered regulation of KA versus 3HK).

### 2.4. Statistics

Analysis of variance (ANOVA) was employed to check differences in continuous variables among diagnostic groups. We used analysis of contingence tables (X^2^-tests) or used the nominal by nominal Ψ measure to check associations between sets of nominal variables. P-correction for false discovery rate (FDR) was used to adjust for multiple comparisons [28]. Bivariate correlations were assessed using Pearson’s product moment correlation coefficients or Spearman’s rank order correlation coefficients. We employed multivariate general linear model (GLM) analysis to delineate the effects of explanatory variables (cytokine levels) on dependent variables (TRYCAT ratios, PHEMN and DAPS symptoms, CERAD tests) while adjusting for age, gender, education and BMI. In case the multivariate analysis yielded significant results we used tests for between-subject effects to assess the univariate effects of significant explanatory variables on the dependent variables. Consequently, model-generated estimated marginal means were computed. We also employed hierarchical multiple regression analysis to delineate the significant explanatory variables (e.g. cytokine levels) predicting CERAD tests, PHEMN and DAPS as dependent variables. All regression analyses were checked for multicollinearity. Exploratory principal component (PC) factor analysis was performed and the PC scores were used in consequent analyses. Adequacy of the correlation matrix was assessed using the Kaiser-Meyer-Olkin (KMO) test and Bartlett’s statistics. To classify participants into relevant biological clusters we used K-means clustering analysis performed on the cytokine/chemokine and TRYCAT data with the main goal to delineate a biological homogeneous class with highly similar biological features. In order to interpret the data structure of the clusters we used ANOVAs, multivariate GLM analysis and binary regression analysis to check differences in clinical and cognitive features between the cluster-analytically generated classes. All statistical analyses were conducted using IBM SPSS windows version 22. Tests were 2-tailed and a p-value of 0.05 was used for statistical significance.

We used multilayer perceptron (MLP) Neural Network analyses (SPSS 22) to decipher the more complex nonlinear relationships between different input variables predicting output variables. The input layer of the automatic feedforward architecture model comprised cytokines, TRYCATs and CERAD tests (with age, sex, BMI and education) and the output levels contained MNP and all other subjects. The relative number of cases assigned to the training (to estimate the network parameters), testing (to prevent overtraining) and holdout (to evaluate the final network) sets were 7, 3 and 5 respectively. The stopping rule was one consecutive step without further decrease in error terms. The automatic architecture model was trained using one or two hidden layers with a variable number of nodes (2-7). We computed error and relative error as well as the (relative) importance of all input variables in sensitivity analyses.

To examine the causal links among cytokine levels, TRYCAT data, neurocognitive deficits and symptom dimensions we used SmartPLS (Partial Least Squares) path modeling analysis [29]. SmartPLS is a structural equation modeling procedure which employs pathway modeling performed on indicator variables or latent constructs extracted from a set of indicator variables. The final response variables were the DAPS dimension (based on HAMA, HAMD and FFs scores), psychotic symptoms (psychosis, hostility, excitement and mannerism scores) and negative symptoms (PANNSneg and SANS). These values were entered as indicator variables in a reflective model. Cytokines/chemokines, the three TRYCAT ratios and CERAD tests were entered as indicators of the latent constructs “immune activation”, “TRYCAT pathway patterning” and “memory deficits”, respectively. Other variables (age, sex, education) were entered as single indicators. TRYCATs and CERAD tests were used as mediators of the effects of immune activation on symptoms. PLS path analysis was only performed when the model and latent constructs complied with specific quality criteria: a) overall quality of the model as assessed with SRMR < 0.08; b) latent constructs should have a good reliability and discriminant validity as indicated by Cronbach’s alpha > 0.7, composite reliability > 0.7 and average variance extracted (AVE) > 0.500; c) indicators of latent constructs should have factor loadings > 0.500 and p values < 0.001; and d) construct crossvalidated redundancies and communalities should be adequate [29]. Subsequently we interpreted path coefficients with exact p-values, total effects, total indirect and specific indirect effects.

## 3. Results

### 3.1. Cluster analysis

Using the K-means method we examined two-cluster solutions using the 4 cytokines and 3 TRYCAT ratios as variables. The clustering method split the data set into 2 classes with 57 (cluster 1) and 62 (cluster 2) subjects. **Table 1** shows the clinical and biomarkers features of both clusters. Firstly, cluster 2 showed higher cytokine/chemokine (all except eotaxin: after FDR p=0.0617) and TRYCAT levels as compared with cluster 1 and is therefore a biologically defined cluster named CYTOTRY cluster, while cluster 1 is named the normal cluster. **Figure 1** shows the z transformed values of the biomarker data in both clusters. Table 1 shows also the socio-demographic features of both clusters: the CYTOTRY cluster showed more males, more single-divorced cases and a lower number of employed participants. The CYTOTRY cluster was also characterized (after FDR) by a greater number of patients with a family history of psychosis, a greater number of psychoses and more hospitalizations and suicidal behavior. There was also a strong association between the cluster-analysis generated biological classes and the diagnostic groups. Thus, more cases allocated to the CYTOTRY cluster had a diagnosis of schizophrenia and especially MNP as compared with the biologically normal cluster, which comprised more controls.

**Figure 1.**
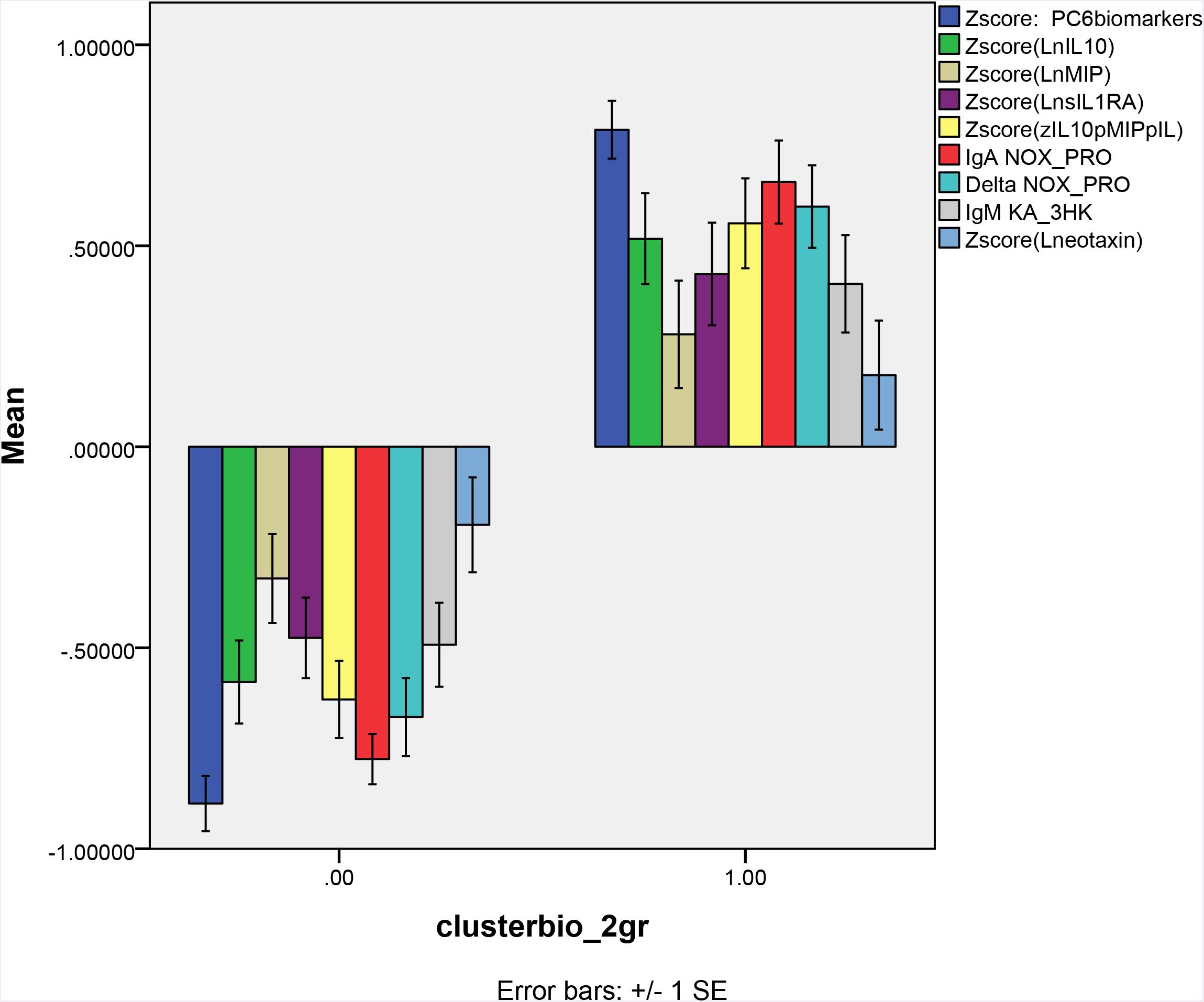
Biomarker measurements in two study groups, namely cluster (1): cluster with increased biomarkers; and cluster (0): normal biomarker values. IL-10: interleukin-10; MIP: macrophage inflammatory protein-1; sIL-1RA: soluble interleukin-1 receptor antagonist; zIL10pMIPpIL: z unit composite score computed as z value of IL-10 (zIL-10) + zMIP-1 + zsIL-1RA; IgA NOX_PRO: computed as sum of z scores of QA (zQA) + zPA + zXA – zAA – zKA (index of increased noxious potential); DelNOX_PRO: computed as IgA (zQA + zPA + zXA +z3HK – zAA – zKA) – zIgM (zQA + zPA + zXA +3HK – zAA – zKA) (a more comprehensive index of increased noxious potential); IgMKA_HK: computed as zIgM KA – z3HK (index of lowered regulation of KA versus 3HK); QA: quinolinic acid; PA: picolinic acid; XA: xanthurenic acid; AA: anthranilic acid; KA: kynurenic acid; 3HK: 3-hydroxy-kynurenine; PC6biomarkers: the first principal components extracted from the 6 above-mentioned immune and TRYCAT markers.

**Table 1.**
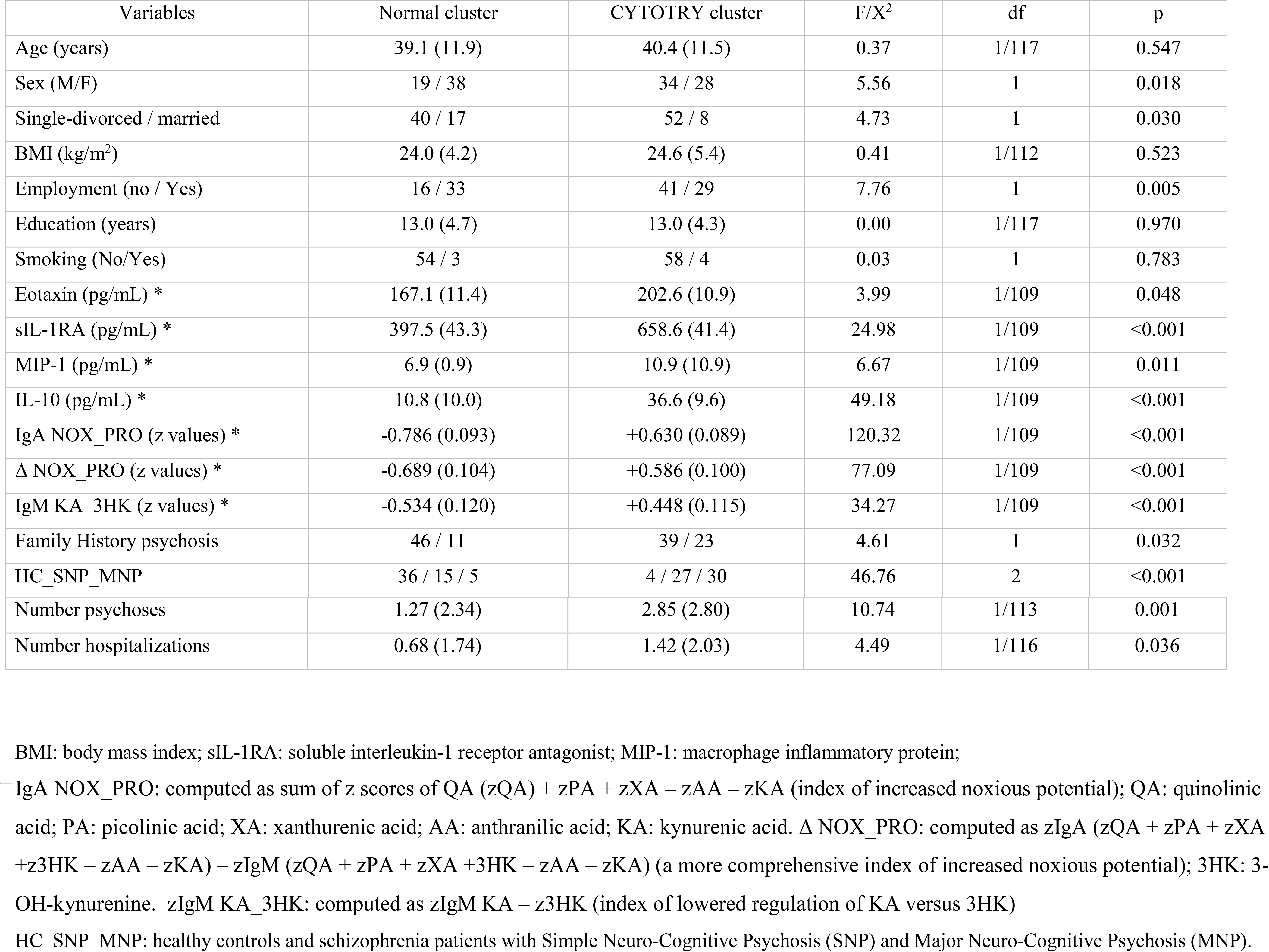
Socio-demographic, clinical and biological data in participants divided according to results of cluster analysis into those with disorders in cytokines/chemokines and tryptophan catabolite (TRYCAT) levels (CYTOTRY cluster).

### 3.2. Factor analysis

The intercorrelation matrices among cytokines and TRYCAT ratios show that (without FDR correction) IL-10 is significantly associated with MIP (r=0.416, p<0.001, all n=120), sIL-1RA (r=0.335, p<0.001), eotaxin (r=0.316, p<0.001), IgA NOX_PRO (r=0.441, p<0.001), Δ NOX_PRO (r=0.302, p=0.001, n=119) and IgM KA_3HK (r=0.344, p<0.001). sIL-1RA is significantly correlated with MIP (r=0.351, p<0.001), IgA NOX_PRO (r=0.305, p=0.001) and Δ NOX_PRO (r=0.244, p=0.007). MIP is significantly associated with IgA NOX_PRO (r=0.232, p=0.011) and Δ NOX_PRO (r=0.215, p=0.019). IgA NOX_PRO was significantly associated with Δ NOX_PRO (r=0.745, p<0.001) and IgM KA_3HK (r=0.408, p<0.001), while NOX_PRO was significantly associated with IgM KA_3HK (r=0.438, p<0.001).

Examining the factor structure of the 7 biomarkers using exploratory factor analysis (PC method) showed that MIP (0.428), sIL-1RA (0.551), IL-10 (0.687), IgA NOX_PRO (0.791), Δ NOX_PRO (0.760) and IgM KA_3HK (0.520) loaded on the first PC and that eotaxin (0.285) did not significantly load on this PC (this first PC explained 35.9% of the total variance). Factor analysis performed on 6 biomarker data without eotaxin showed a better solution whereby 41.1% of the variance was explained (KMO=0.612 which is moderate; Bartlett’s test of sphericity: X^2^=175.15, df=15, p<0.001). MIP (0.427), sIL-1RA (0.542), IL-10 (0.657), IgA NOX_PRO (0.808), Δ NOX_PRO (0.782) and IgM KA_3HK (0.540) loaded on the first PC. Moreover, the latent vector extracted from these 6 biomarkers shows a good composite reliability (0.796). Consequently, we have computed the factor score on this first PC and used this score in subsequent analyses (named: PC 6 biomarkers). This score thus reflects a severity score of immune-TRYCAT disorders (all without eotaxin). In addition, we have computed a z unit weighted composite score based on z values of IL-10 (zIL10) levels + zsIL-1RA + zMIP (named: zIL1RA+MIP+IL10). This score reflects a general immune activation associated with M1 and Th-2/Treg activities.

### 3.3. Predicting MNP and SNP using immune variables using logistic regression analysis

**Table 2** shows the results of hierarchical binary logistic regression analyses with SCZ or MNP as dependent variables and controls or SNP as reference groups. Regression #1 shows that SCZ (versus HC) was best predicted by CYTOTRY cluster, eotaxin and IL-10 combined (X^2^=88.51, df=3, p<0.001, Nagelkerke=0.728); 88.2% of all cases were correctly classified with a sensitivity of 91.1% and a specificity of 82.5%. Regression #2 shows that MNP (versus SNP) was best predicted by increased scores on the PC 6 biomarkers and lowered BMI (X^2^=23.71, df=2, p<0.001, Nagelkerke=0.375; 79.2% of all cases were correctly classified with sensitivity of 78.7% and specificity 79.5%). Regression #3 shows that MNP (versus controls) was best predicted by PC 6 biomarkers and eotaxin (X^2^=68.22, df=2, p<0.001, Nagelkerke=0.812; 91.8% of all cases were correctly classified with sensitivity of 90.9% and specificity 92.5%). Regression #4 shows that SNP (versus controls) was best predicted by increased levels of IL-10, sIL-1RA and eotaxin (X^2^=62.81, df=3, p<0.001, Nagelkerke=0.730; 86.1% of all cases were correctly classified with sensitivity of 87.2% and specificity 85.0%).

**Table 2.**
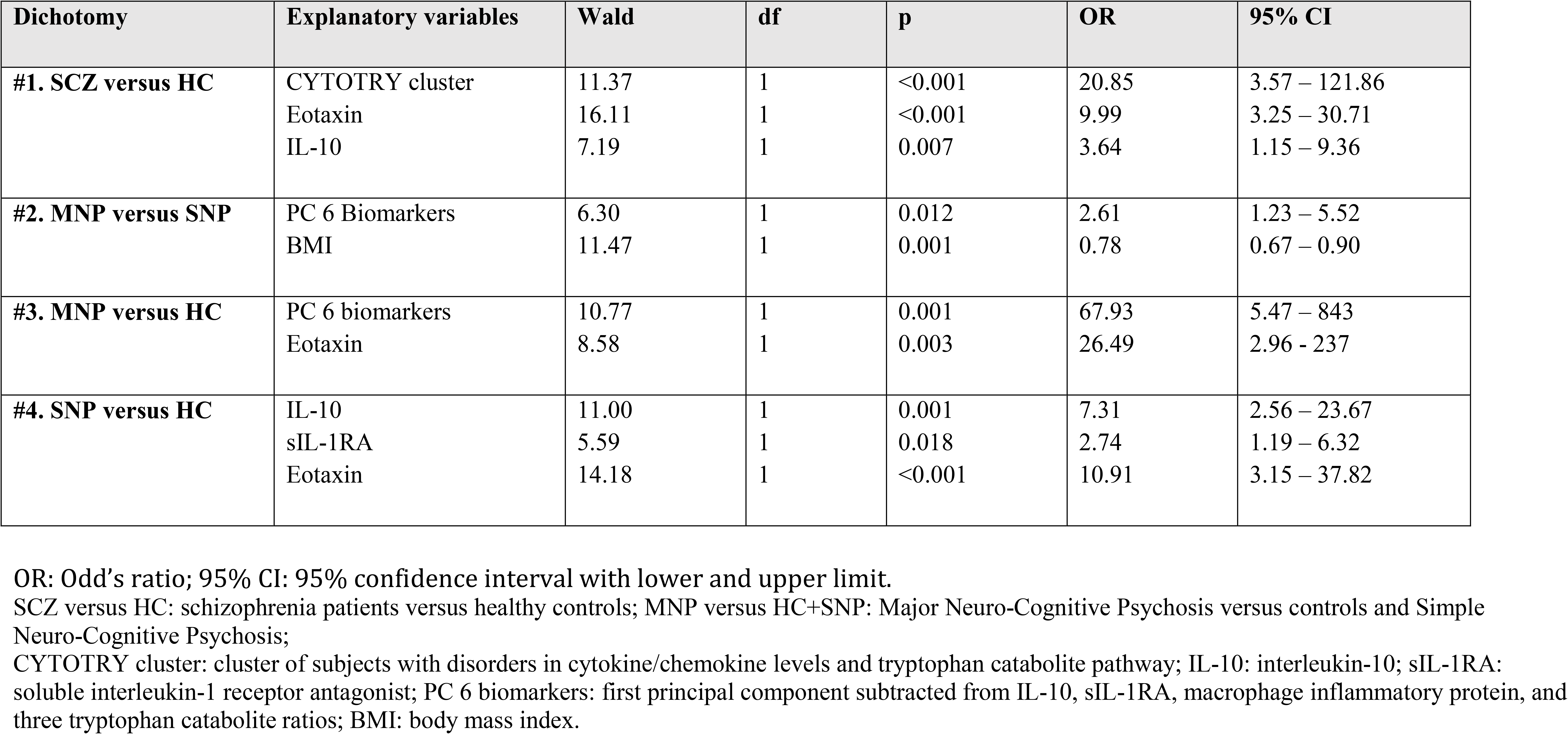
Results of binary logistic regression analyses with diagnosis of schizophrenia (SCZ) or Major Neuro-Cognitive Psychosis (MNP) as dependent variables

| Dichotomy | Explanatory variables | Wald | df | p | OR | 95% CI |
| --- | --- | --- | --- | --- | --- | --- |
| #1. SCZ versus HC | CYTOTRY cluster | 11.37 | 1 | <0.001 | 20.85 | 3.57 – 121.86 |
|  | Eotaxin | 16.11 | 1 | <0.001 | 9.99 | 3.25 – 30.71 |
|  | IL-10 | 7.19 | 1 | 0.007 | 3.64 | 1.15 – 9.36 |
| #2. MNP versus SNP | PC 6 Biomarkers | 6.30 | 1 | 0.012 | 2.61 | 1.23 – 5.52 |
|  | BMI | 11.47 | 1 | 0.001 | 0.78 | 0.67 – 0.90 |
| #3. MNP versus HC | PC 6 biomarkers | 10.77 | 1 | 0.001 | 67.93 | 5.47 – 843 |
|  | Eotaxin | 8.58 | 1 | 0.003 | 26.49 | 2.96 – 237 |
| #4. SNP versus HC | IL-10 | 11.00 | 1 | 0.001 | 7.31 | 2.56 – 23.67 |
|  | sIL-1RA | 5.59 | 1 | 0.018 | 2.74 | 1.19 – 6.32 |
|  | Eotaxin | 14.18 | 1 | <0.001 | 10.91 | 3.15 – 37.82 |

OR: Odd's ratio; 95% CI: 95% confidence interval with lower and upper limit.
SCZ versus HC: schizophrenia patients versus healthy controls; MNP versus HC+SNP: Major Neuro-Cognitive Psychosis versus controls and Simple Neuro-Cognitive Psychosis;
CYTOTRY cluster: cluster of subjects with disorders in cytokine/chemokine levels and tryptophan catabolite pathway; IL-10: interleukin-10; sIL-1RA: soluble interleukin-1 receptor antagonist; PC 6 biomarkers: first principal component subtracted from IL-10, sIL-1RA, macrophage inflammatory protein, and three tryptophan catabolite ratios; BMI: body mass index.

### 3.4. Predicting MNP and SNP using immune variables in neural network (NN) analysis

Using NN analyses we have examined the predictors of schizophrenia (versus controls) and MNP (versus controls + SNP). Table 3 shows the model summary of the trained Neural Networks as well as the confusion matrices. We trained both models entering 14 units as input variables, namely all 7 biomarkers, CYTOTRY cluster, indices of semantic and episodic memory, age, sex, education and BMI. Automatic architecture training of the network predicting schizophrenia delineated the best model by using 2 hidden layers, with 3 units in the first hidden layer and 2 units in the second hidden layer. The model used hyperbolic tangent as activation function in the first layer and identity in the output layer and used a sum of squares error term. NN #1 in Table 3 shows that the sum of squares error term was minimized during the training, indicating that the model learnt to generalize from the trend. The percentage of incorrect classifications was lower in the testing and holdout samples than in the training sample indicating that the model is not overtrained (overfitted) and will generalize well. The AUC ROC curve was 0.985 and the confusion matrix in the holdout sample showed a sensitivity of 100% and specificity of 82.4%. **Figure 2** shows the relevance chart with normalized and relative importances of all input variables in the model. Eotaxin and IgA NOX_PRO ratio were the most important determinants of the predictive power of the model, followed at a distance by IL-10 and Δ NOX_PRO and again at a distance by MIP.

**Figure 2.**
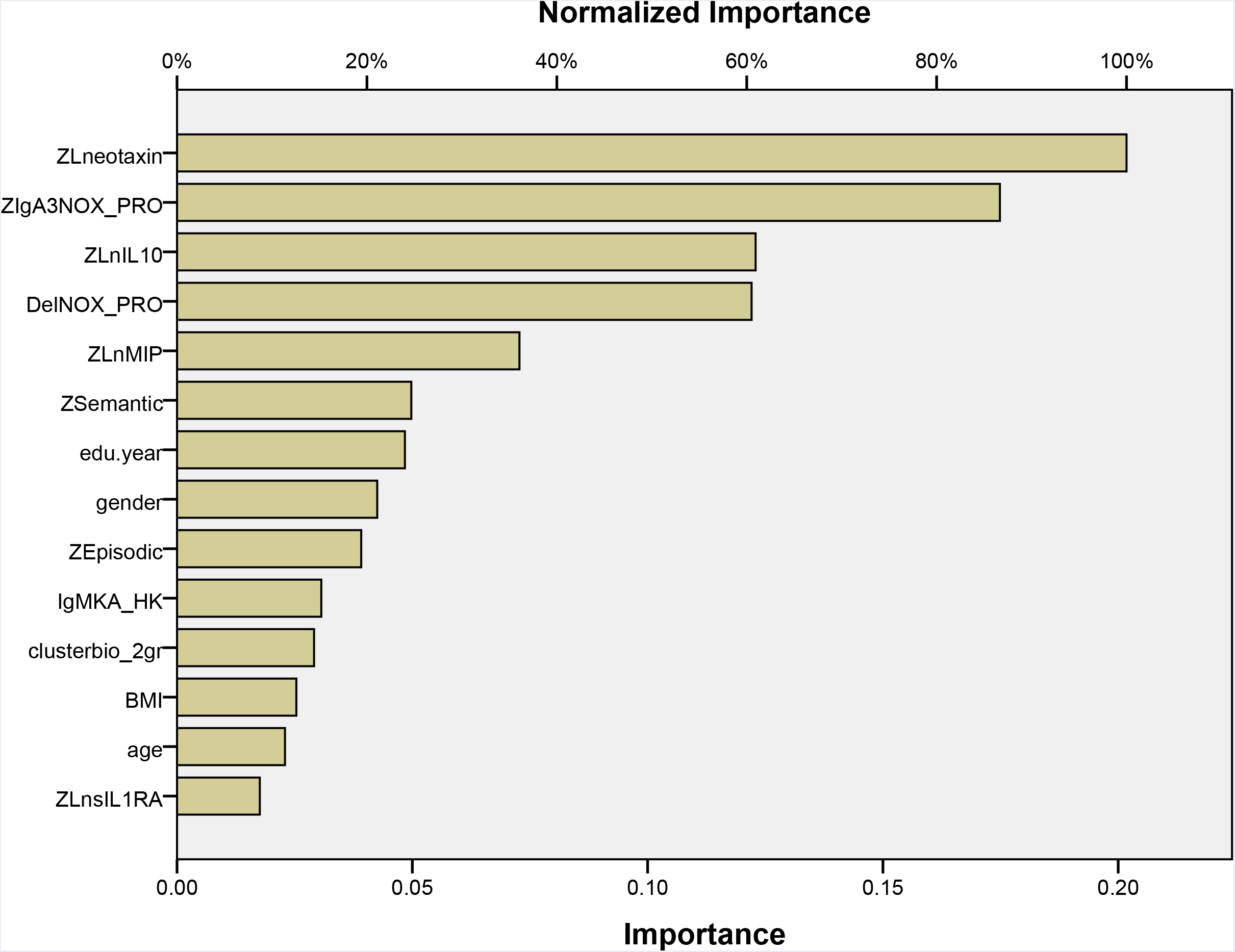
Results of Neural network analysis with schizophrenia versus controls as output variables. See legends to Table 1 for explanation on biomarkers used. Cluterbio_2gr: the two clusters as shown in Figure 1 entered as dummy variable; ZSemantic: Principal component extracted from semantic memory tests; ZEpisodic: Principal component extracted from episodic memory tests; BMI: body mass index.

**Table 3.** Results of Neural network analysis with schizophrenia (SCZ) or Major Neuro-Cognitive Psychosis (MNP) as output variables and cytokine levels, tryptophan catabolites, cognitive data and socio-demographic data as input variables. Shown are the model summary, the partitioned confusion matrix (predicted) and the area under the ROC curve (AUC ROC).

| Dichotomy | Samples | Sum of squares error term | % incorrect | Observed | Predicted | AUC ROC |
| --- | --- | --- | --- | --- | --- | --- |
| #1. SCZ versus HC | Training | 3.571 | 14.5% | HC<br>SCZ | 53.8%<br>95.2% | 0.985 |
|  | Testing | 1.582 | 10.0% | HC<br>SCZ | 80.0%<br>100.0% |  |
|  | Holdout | - | 7.9% | HC<br>SCZ | 82.4%<br>100.0% |  |
| #2. MNP versus all others | Training | 5.862 | 13.5% | HC+SNP<br>MNP | 91.2%<br>77.8% | 0.928 |
|  | Testing | 0.732 | 0.0% | HC+SNP<br>MNP | 100%<br>100% |  |
|  | Holdout | - | 12.5% | HC+SNP<br>MCP | 96.7%<br>60.0% |  |

SCZ versus HC: schizophrenia patients versus healthy controls; MNP versus HC+SNP: Major Neuro-Cognitive Psychosis versus controls and Simple Neuro-Cognitive Psychosis

**Table 4.**
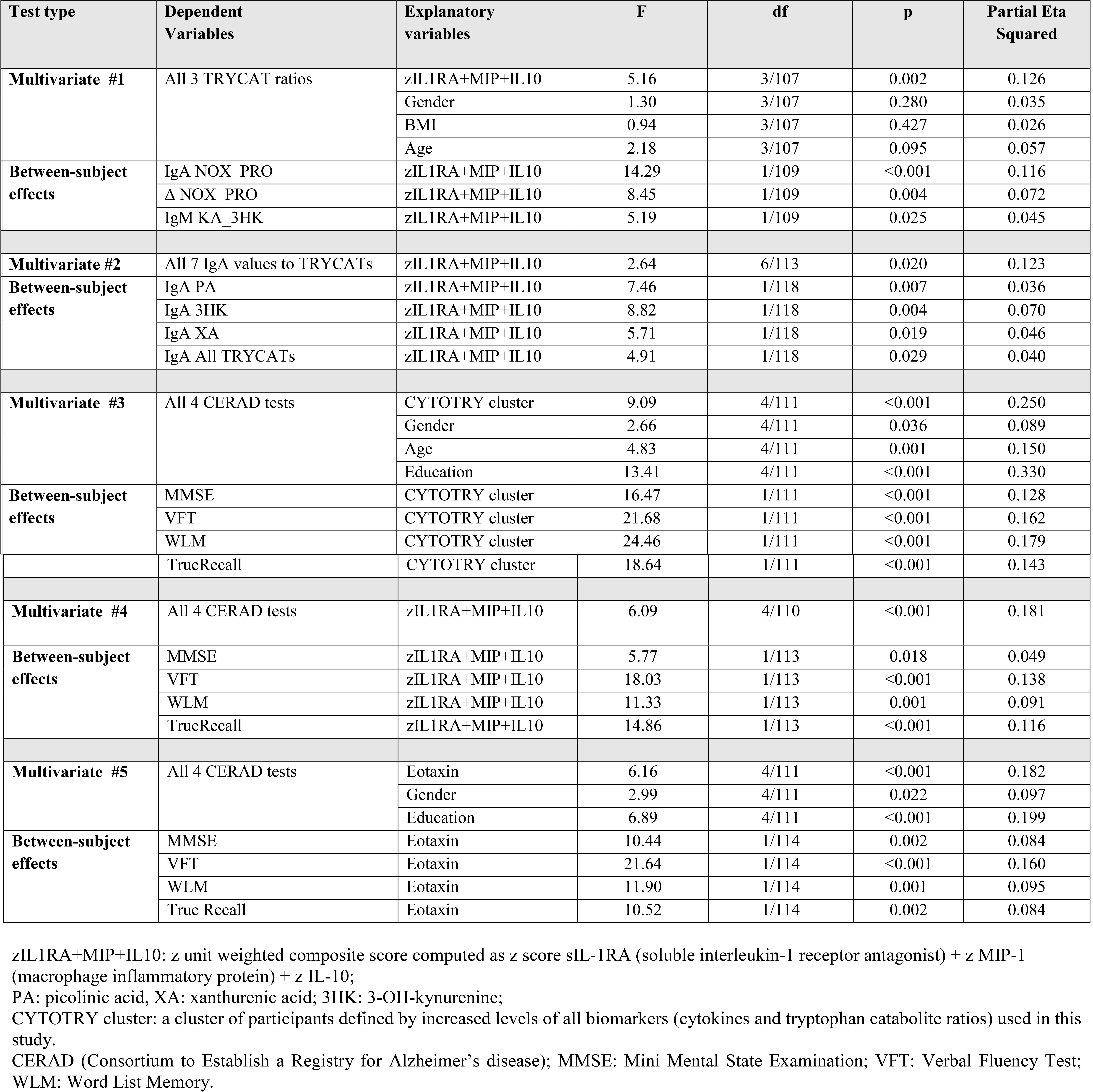
Results of multivariate GLM analyses with tryptophan catabolite (TRYCATs) ratios as dependent variables and cytokine/chemokine levels or the CYTOTRY (cytokine-tryptophan catabolite) cluster as explanatory variablesz

NN #2 in Table 3 shows a second analysis with MNP versus all other subjects as output variables. This model was trained using 2 hidden layers, with 2 units in the first and second hidden layers and using hyperbolic tangent as activation function in the first layer and identity in the output layer. Table 3 shows that the sum of squares error term was minimized during the training and that the percentage of incorrect classifications was lower in the testing and holdout samples than in the training sample, indicating that the model is not overfitted and may generalize well. **Figure 3** shows the relevance chart with the importances of the input variables. Education, BMI, IgM KA_3HK and the CYTOTRY cluster were the dominant determinants of the predictive power of the model, followed at a distance by episodic memory and eotaxin.

**Figure 3.**
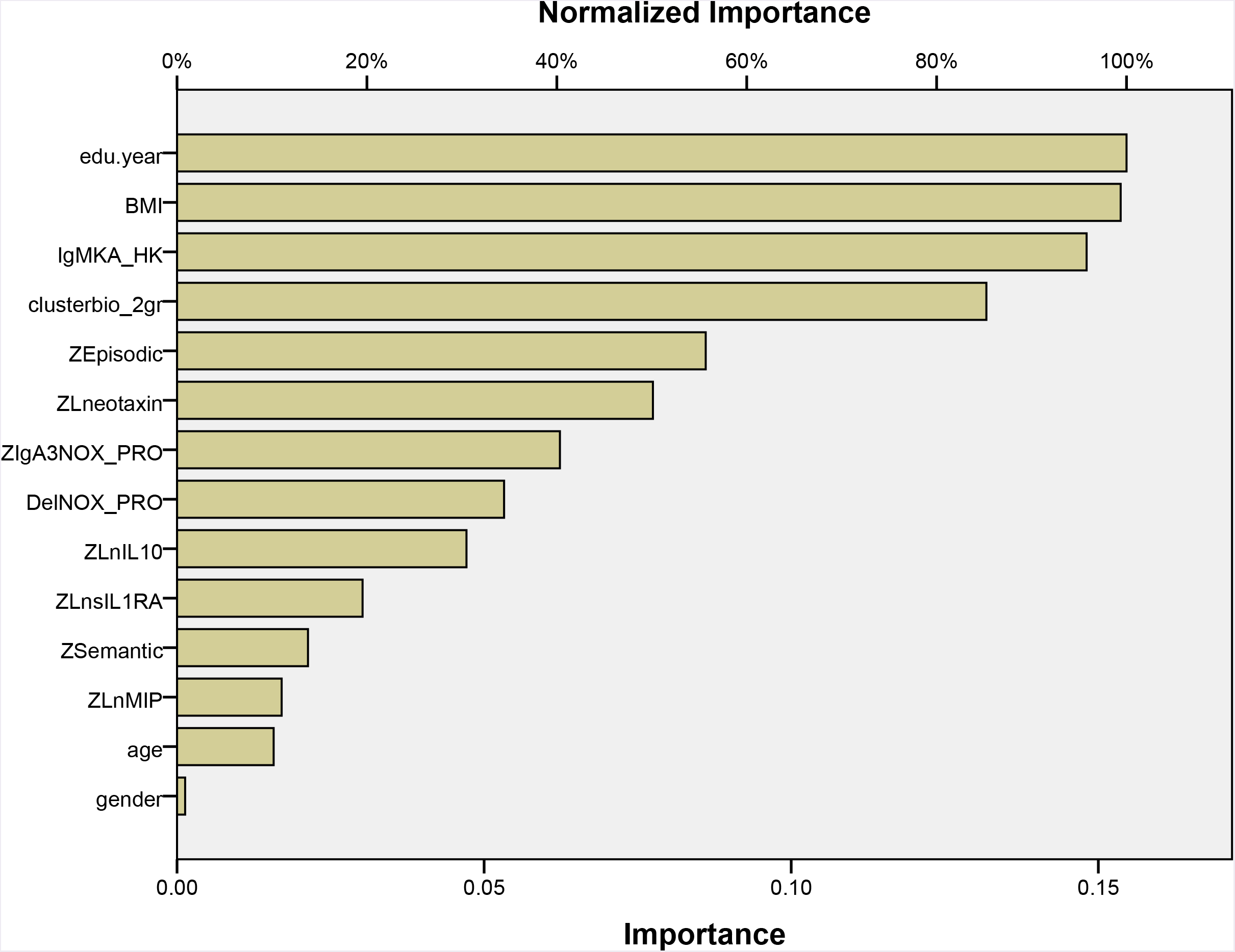
Results of Neural network analysis with Major Neuro-Cognitive Psychosis (MNP) versus Simple Neuro-Cognitive Psychosis and controls as output variables.

### 3.5. Predicting TRYCAT patterning by indices of immune activation

**Table 4** regression #1 shows the results of multivariate GLM analysis with 3 TRYCAT ratios as dependent variables and zIL1RA+MIP+IL10 score as explanatory variable while adjusting for gender, age and BMI (these three extraneous variables were not significant). There was a significant effect of the immune activation index on TRYCATs with an explained variance around 12.6%. Tests for between-subjects effects showed significant effects of the immune activation index on all three ratios with a stronger effect size on IgA NOX_PRO. We have also examined the effects of immune activation on IgA responses directed to all 6 TRYCATs and their sum (regression #2 in Table 4) and found that there were significant effects of immune activation increasing PA, 3HK and XA levels as well as the sum of all TRYCATs. Nevertheless, there was no significant impact of eotaxin on the TRYCAT levels (results of multivariate GLM analysis: F=0.29, df=3/115, p=0.831).

### 3.6. Predicting cognitive impairments by immune variables

We have examined the effects CYTOTRY cluster, zIL1RA+MIP+IL10 or eotaxin (entered as explanatory variables) on CERAD tests (MMSE, VFT, WLM, TrueRecall; entered as dependent variables while adjusting for age, sex and education). Regression #3 shows that around 25.0% of the variance in CERAD tests could be explained by the regression on CYTOTRY cluster and that all 4 CERAD tests were strongly predicted by the CYTOTRY cluster. Regression #4 shows that the immune index zIL1RA+MIP+IL10 explained around 18.1% in the CERAD tests and that there were strong associations between immune activation and VFT, WLM and TrueRecall. Regression # 5 shows that eotaxin also predicted cognitive functioning whereby 18.2% of the variance in the CERAD tests was explained by the regression on eotaxin. In addition, eotaxin significantly impacted all 4 CERAD tests and especially VFT.

### 3.7. Prediction of symptom dimensions using cytokines/chemokines

In order to examine the effects of immune activation on schizophrenia symptom dimensions we examined the effects of the CYTOTRY cluster, zIL1RA+MIP+IL10 or eotaxin as explanatory variables on the 8 symptom dimensions (entered as dependent variables in multivariate GLM analysis while adjusting for age, sex and education). **Table 5**, Regression #1 shows that 42.8% of the variance in the 8 symptom dimensions was explained by the CYTOTRY cluster with a very strong impact on SANS, HAMD and FF, a strong effect on excitation, HAMA and psychosis and more moderate effects on mannerism, but not hostility. Regression #2 shows that 19.2% of the variance in the 8 symptom dimensions is explained by the zIL1RA+MIP+IL10 score which yielded significant effects on all dimensions (including hostility) with particularly strong effects on HAMD, FF and SANS. Regression #3 shows that 19.8% of the variance in the symptoms is explained by the regression on eotaxin, which strongly predict HAMD, FF and SANS with moderate effects on psychosis, excitation and mannerism (but no effects on hostility and HAMA).

**Table 5.**
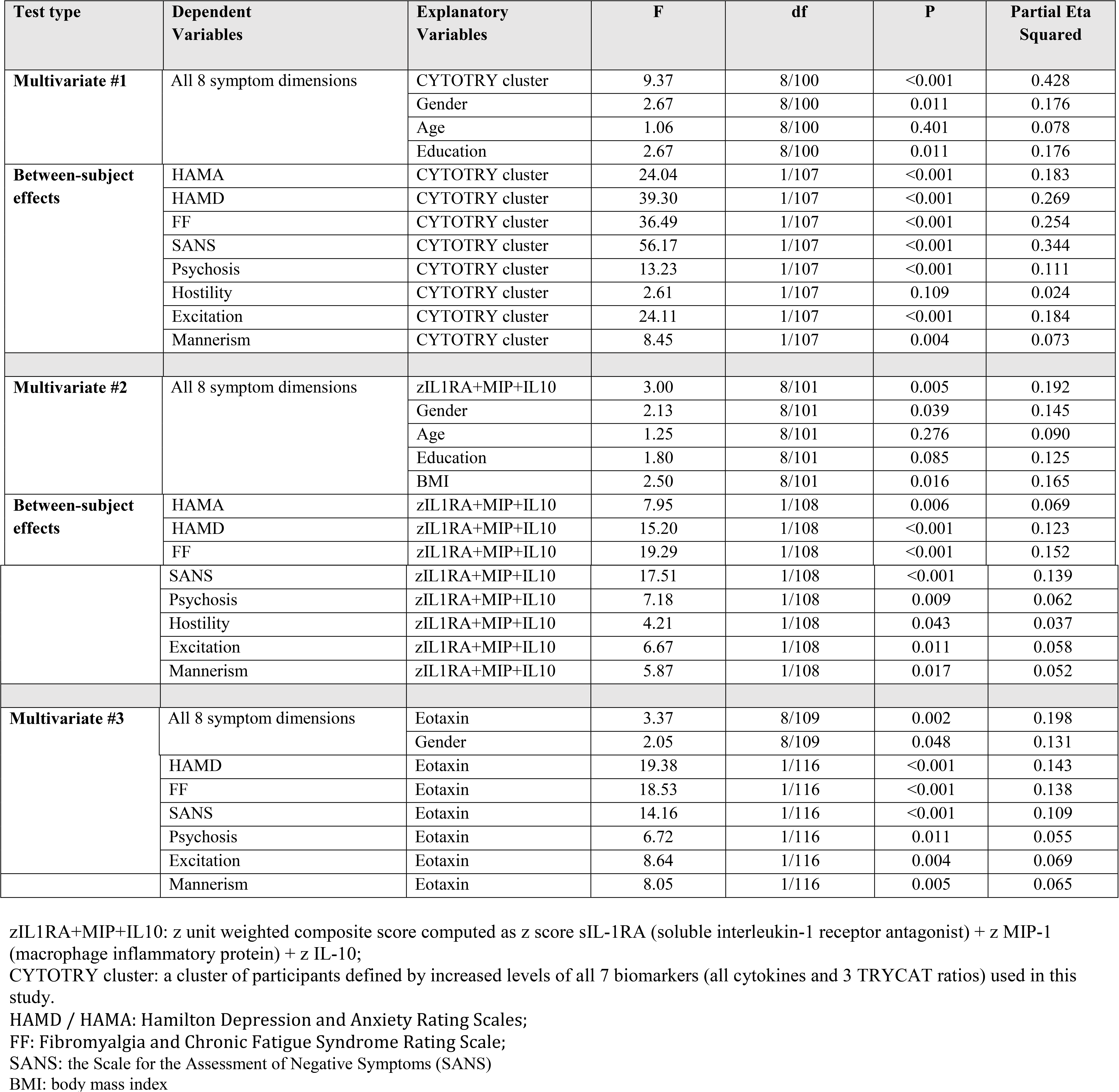
Results of multivariate GLM analyses with clinical symptom dimensions as dependent variables and cytokines/chemokines and the CYTOTRY (cytokines-tryptophan catabolite) cluster as explanatory variables

**Table 6.**
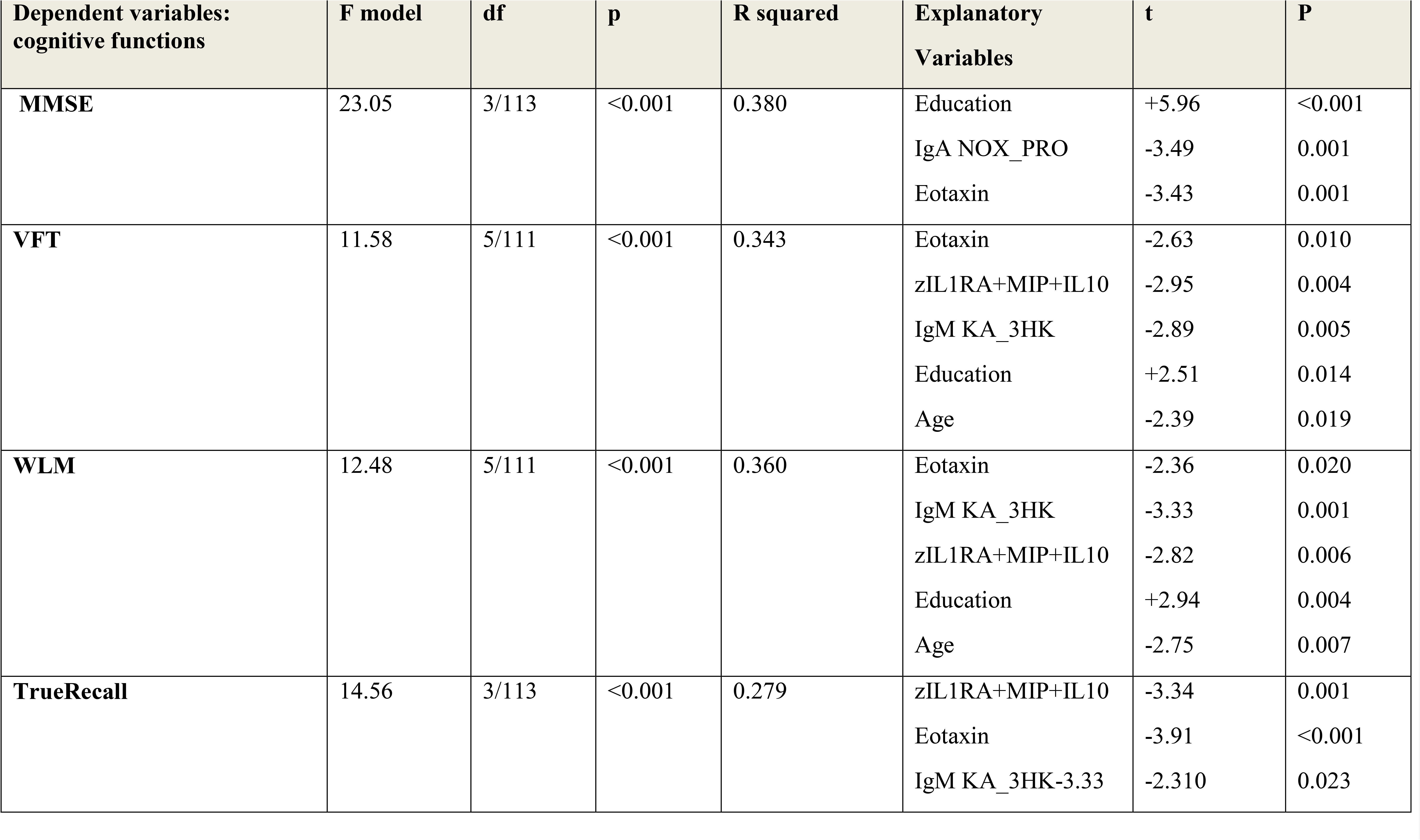
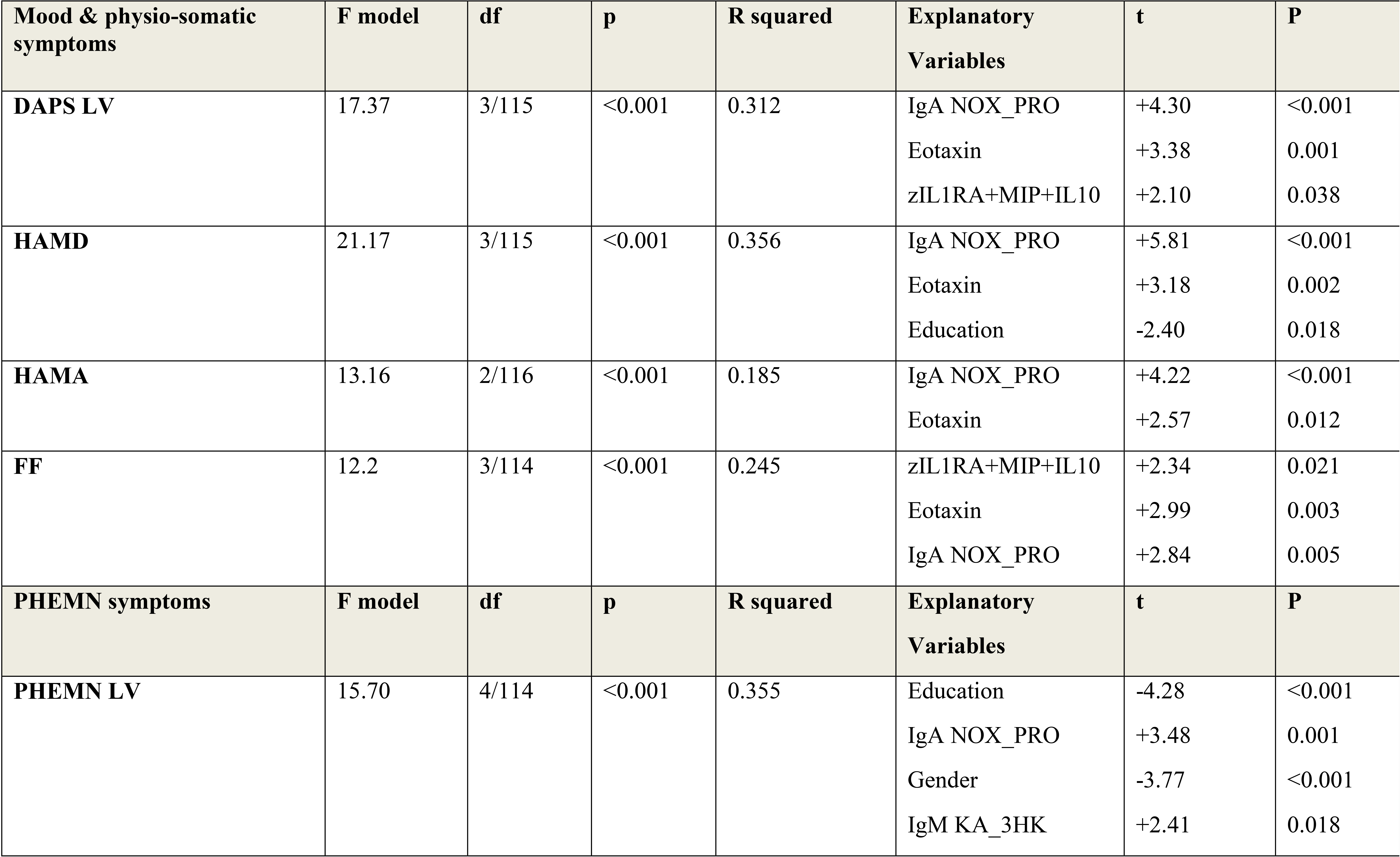
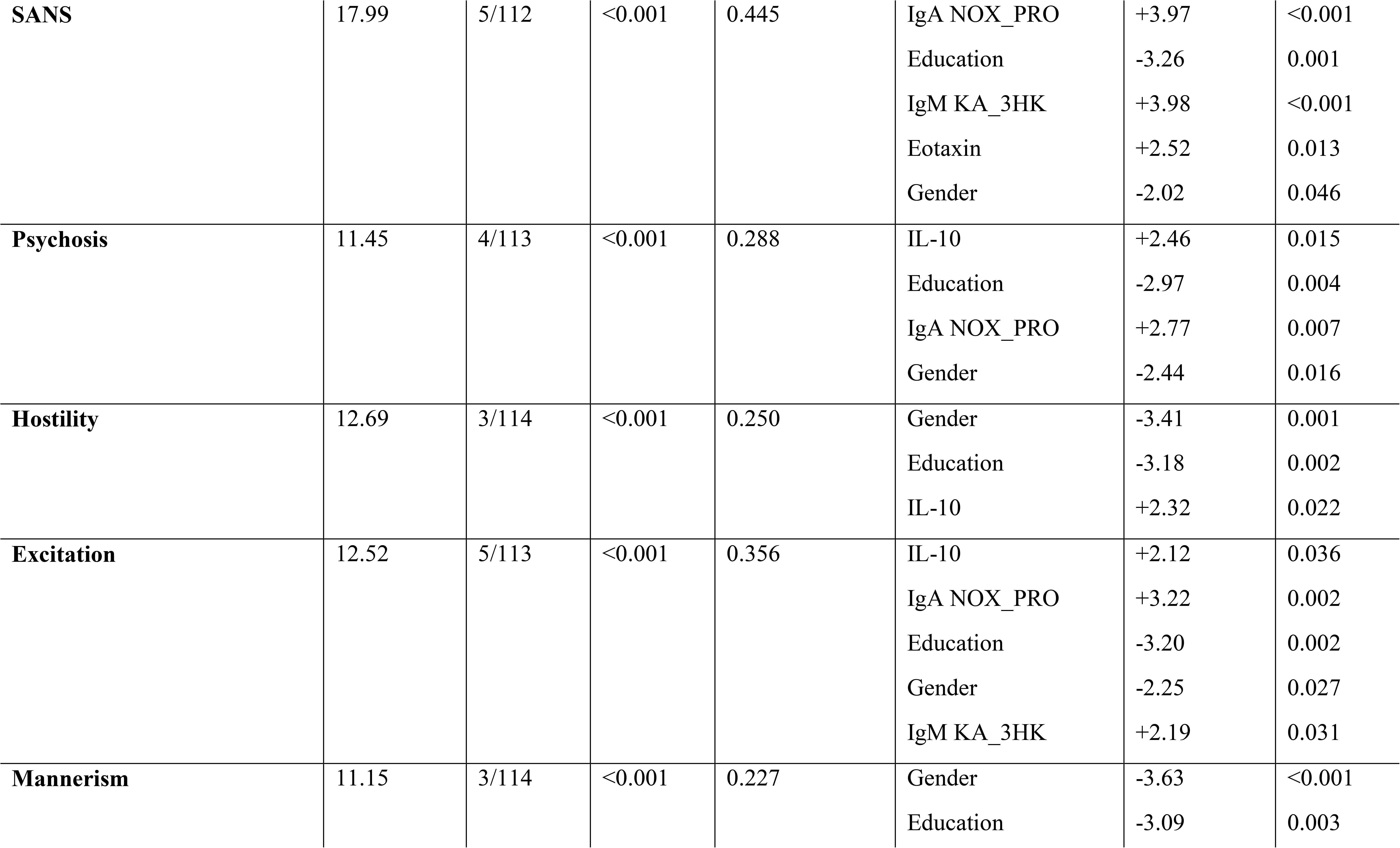
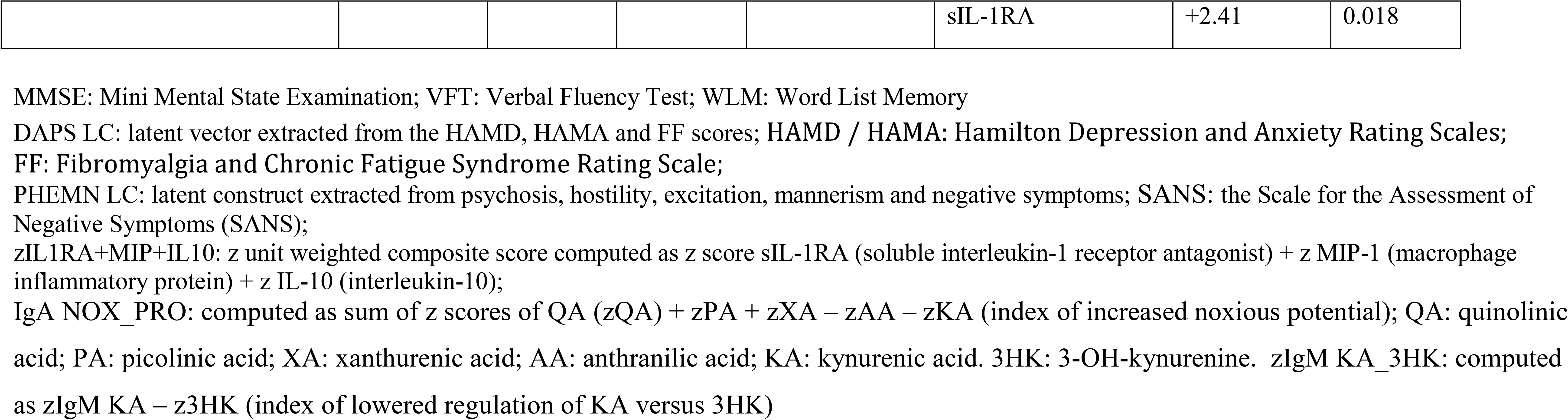
Results of hierarchical regression analyses with different cognitive tests and symptom dimensions as dependent variablesMMSE: Mini Mental State Examination; VFT: Verbal Fluency Test; WLM: Word List Memory

### 3.8. Prediction of cognitive functions and symptom dimensions using immune and TRYCAT biomarkers

In order, to delineate the best prediction of the different CERAD and symptom dimensions using cytokines/chemokines and TRYCAT data as explanatory variables we performed hierarchical multiple regression analyses. MMSE was best predicted by education, IgA NOX_PRO ratio and eotaxin (38.0% of the variance). VFT was best predicted by eotaxin, zIL1RA+MI+IL10, IgM KA_3HK, education and age (34.3% of the variance). 36.0% of the variance in WLM was explained by the regression on eotaxin, IgM KA_3HK, zIL1RA+MI+IL10, education and age. 27.9% of the variance in TrueRecall was predicted by zIL1RA+MI+IL10, eotaxin and IgM KA_3HK.

The DAPS LV (latent vector) was best predicted by IgA NOX_PRO, eotaxin and zIL1RA+MI+IL10 (31.2% of the variance). The same variables explained 24.5% of the variance in FF score. IgA NOX_PRO and eotaxin combined with education explained 35.6% of the variance in HAMD, while IgA NOX_PRO and eotaxin explained 18.5% of the variance in HAMA.

The PHEMN LV was best predicted by IgA NOX_PRO, IgM KA_3HK, lower education and male sex (35.5% of the variance). A large part of the variance in the SANS (44.5%) was explained by IgA NOX_PRO, IgM KA_3HK, eotaxin, lower education and male sex. Psychosis was strongly predicted by IL-10, IgA NOX_PRO, lower education and male sex, and hostility by IL-10, male sex and lower education. A larger part of the variance in excitation (35.6%) was explained by the regression on IL-10, IgA NOX_PRO, IgM KA_3HK, lower education and male sex. Finally, 22.7% of the variance in mannerism could be explained by sIL-1RA levels, male sex and lower education.

### 3.9. Prediction of symptom dimensions by immune effects on TRYCATs and cognition: results of PLS path modeling

In **Figure 4** we examine the causal links between the immune activation index and eotaxin and TRYCATs and the effects of these input variables on three symptom dimensions, namely DAPS, PHEM and negative symptoms. Negative symptoms were entered separately from PHEM to examine differential effects of the input variables on both dimensions (although negative symptoms and PHEM belong to the same PHEMN dimension). The overall model fit was good (SRMR=0.075), and the construct reliability and discriminant validity of all latent constructs were good to excellent (all composite reliability > 0.7 and average variance extracted > 0.5). The construct crossvalidated redundancies and communalities were adequate. Finally, all factor loadings of the 4 constructs showed loadings > 0.55, indicating that the convergent validity is adequate. Figure 6 shows the path coefficients with exact p values. We found that 35.8% of the variance in the PHEM LV was explained by the regression on TRYCATs, education and sex, while there were no effects of eotaxin and cytokines. 45.8% of the variance in negative symptoms was explained by eotaxin, TRYCATs, and education. 34.1% of the variance in DAPS dimension was explained by eotaxin, TRYCATs and education. 30.4% of the variance in eotaxin was associated with age and sex (and cytokines were nearly significant). Finally, 25.8% of the variance in TRYCATs was explained by immune activation. There were specific indirect effects of cytokines on DAPS (t=+2.67, p=0.008), negative symptoms (t=+3.22, p=0.001) and PHEM (t=+2.57, p=0.011) which were mediated by TRYCATs. There were highly significant total effects of cytokines on DAPS (t=+3.34, p=0.001), negative symptoms (t=+4.03, p<0.001) and PHEM (t=+3.08, p=0.002).

**Figure 4.**
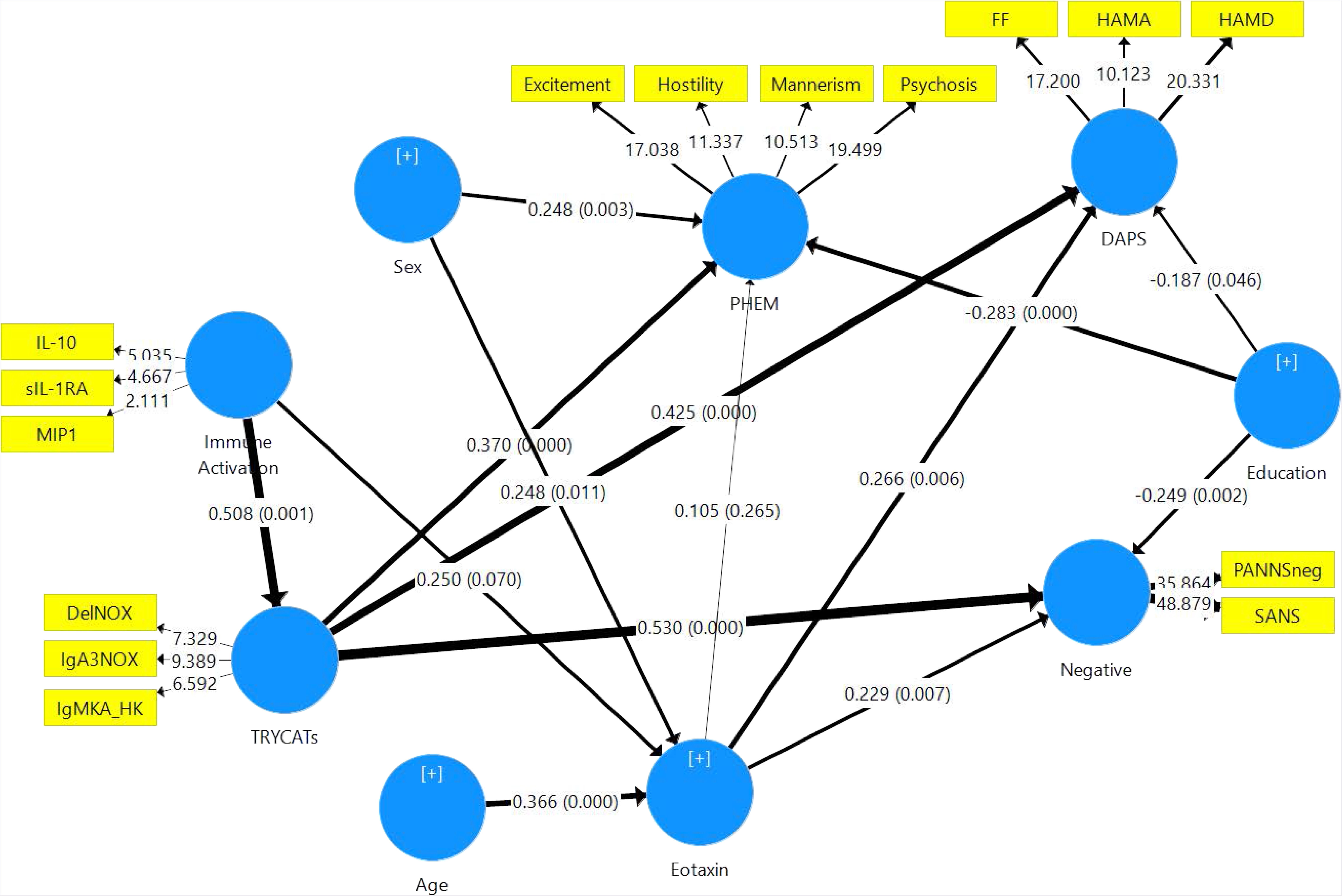
Results of Partial Least Squares (PLS) analysis with DAPS (Depression – Anxiety – Physio-Somatic), PHEM (Psychosis – Hostility – Excitation – Mannerism) and negative symptoms as measured with SANS (Scale for the Assessment of Negative Symptoms) and PANNSnegative (negative subscale of the Positive and Negative Syndrome Scale) as outcome variables. IL-10: interleukin-10; MIP: macrophage inflammatory protein-; sIL-1RA: soluble interleukin-1 receptor antagonist; IgA NOX_PRO: computed as sum of z scores of QA (zQA) + zPA + zXA – zAA – zKA (index of increased noxious potential); DelNOX_PRO: computed as IgA (zQA + zPA + zXA +z3HK – zAA – zKA) – zIgM (zQA + zPA + zXA +3HK – zAA – zKA) (a more comprehensive index of increased noxious potential); IgMKA_HK: computed as zIgM KA – z3HK (index of lowered regulation of KA versus 3HK); QA: quinolinic acid; PA: picolinic acid; XA: xanthurenic acid; AA: anthranilic acid; KA: kynurenic acid; 3HK: 3-hydroxy-kynurenine; HAMD / HAMA: Hamilton Depression and Anxiety Rating Scales; FF: Fibromyalgia and Chronic Fatigue Syndrome Rating Scale.

**Figure 5** shows a similar model but now with inclusion of CERAD tests in order to examine whether the effects of immune activation and cytokines on symptom dimensions are (in part) mediated via cognitive impairments. The model quality data were fairly good with an overall model fit SRMR=0.073, and the construct reliability and discriminant validity of all latent constructs were good to excellent (all composite reliability > 0.7 and average variance extracted > 0.5). This model showed that 51.3% of the variance in PHEM was explained by memory and sex, and 68.6% of the variance in negative symptoms by CERAD tests and TRYCATs. 47.4% of the variance in DAPS was explained by CERAD tests, while 58.8% of the variance in memory tests was explained by eotaxin, TRYCATs and education. There were specific indirect effects of immune activation on DAPS (t=+2.50, p=0.013), negative symptoms (t=+2.58, p=0.010) and PHEM (t=+2.33, p=0.020), which were mediated by effects of TRYCAT on CERAD tests, and of immune activation on negative symptoms mediated by TRYCATs (t=+2.46, p=0.014). There were also specific indirect effects of immune activation on memory (t=2.61 p=0.009) mediated by TRYCATs. In addition, there were significant indirect effects of eotaxin on DAPS (t=+3.14, p=0.002), PHEM (t=+3.16, p=0.002) and negative symptoms (t=+3.41, p=0.001), which were mediated by memory deficits.

**Figure 5.**
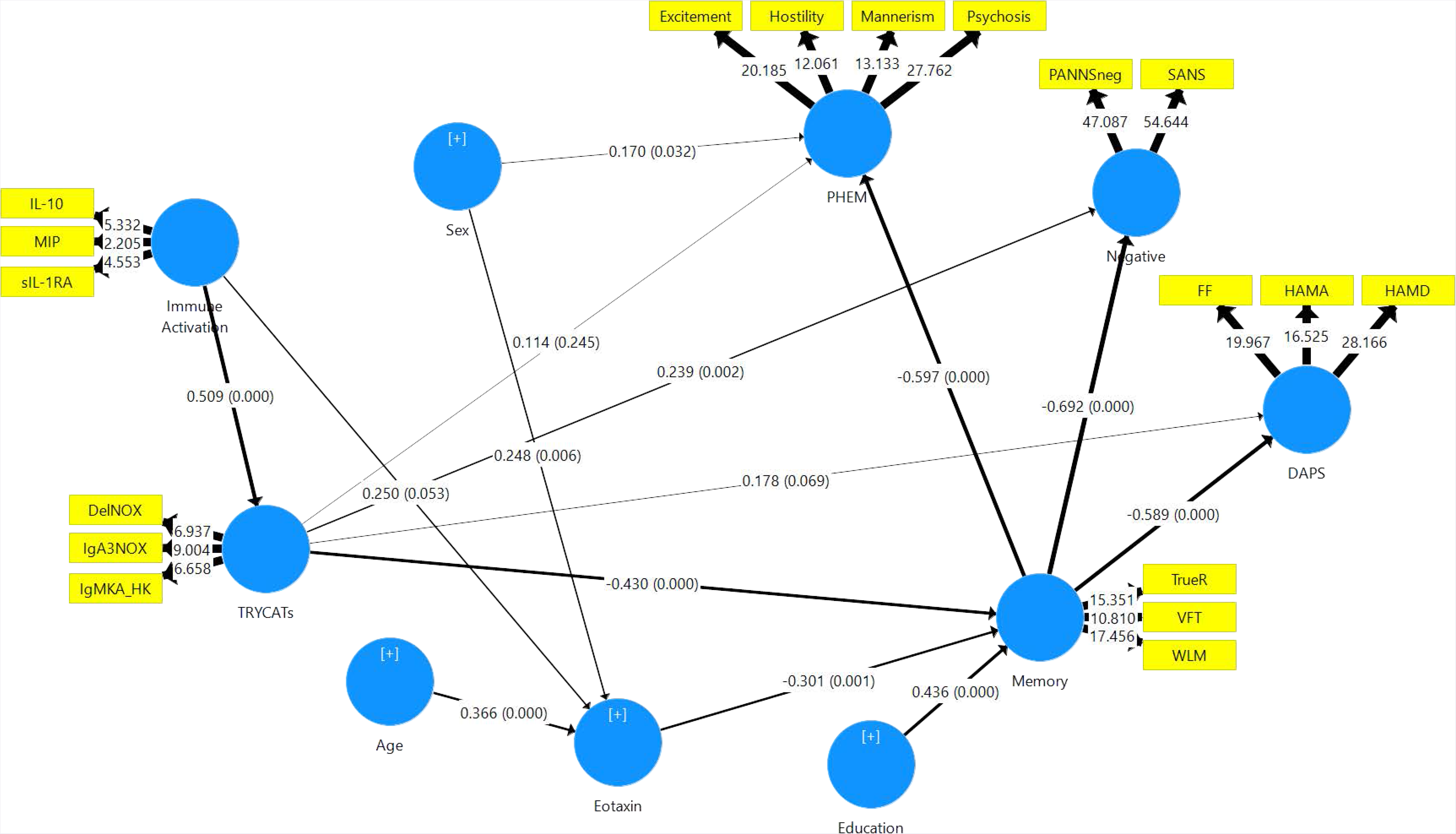
Results of a Partial Least Squares (PLS) Analysis. This PLS model is similar to that shown in Figure 6, but here we have included Memory tests as additional explanatory variables. Memory: as assessed with CERAD (Consortium to Establish a Registry for Alzheimer’s disease) tests, namely WLM: Word List Memory; VFT: Verbal Fluency Test, and TrueR: True Recall.

## 4. Discussion

### 4.1. Immune activation and schizophrenia

The first major finding of this study is that schizophrenia, SNP and MNP are significantly associated with signs of immune activation (increased sIL-1RA, IL-10 and/or MIP-1) and increased eotaxin. Increased sIL-1RA levels indicate M1 activation and increased IL-1β release, while increased IL-10 indicates Treg (Th-2) activation [30]. MIP-1 (CCL3) is a chemokine released by macrophages during an immune response that may activate neutrophils and eosinophils and the secretion of IL-1, IL-6 and TNF-α [31,32]. Eotaxin, a chemotaxin that attracts eosinophils, is induced by Th-2 cytokines, including IL-4, IL-10 and IL-13, and secreted by B cells, eosinophils, endothelial cells, macrophages, fibroblasts, lymphocytes, epithelial cells and chondrocytes [33-36]. As described in the Introduction, there is now evidence that schizophrenia is accompanied by activated M1, Th-1, Th-2 and Treg responses. Also, other authors found increased MIP-1 and eotaxin levels in schizophrenia patients [16,37-40]. In the present study, we found that using eotaxin, IL-10, immune and TRYCAT data yielded a good diagnostic performance for schizophrenia with sensitivity equaling 83.8% and specificity 90.0%. Previously, increased eotaxin levels combined with other immune biomarkers (e.g. soluble TNF receptors and MIP-1) yielded a good diagnostic performance (sensitivity: 70.0% and specificity: 89.4%) for schizophrenia [16]. Importantly, here we observed that eotaxin together with immune/TRYCAT data showed a good diagnostic performance for MNP, namely sensitivity=90.9% and specificity=92.5%. These data further underscore that MNP is biologically validated as an immune-mediated diagnostic category.

Nevertheless, it should be stressed that the immune index used here, namely zsIL1RA+MIP+IL10, not only indicates M1 (sIL-1RA and MIP-1) and Th-2/Treg (IL-10), but also Th-1 activation. This because M1, Th-1, Th-2 and Treg activation are interrelated phenomena that co-occur in acute phases of schizophrenia, chronic schizophrenia and first episode schizophrenia [15-17]. Thus, in the current study, IL-10 was significantly associated with sIL-1RA, MIP-1 and eotaxin levels. Also, in our previous studies we established that IL-10 (as an indicant of Treg and Th-2 functions) was consistently increased in schizophrenia and exhibited strong associations with M1 and Th-1 cytokines [17]. In addition, while sIL-1RA is a product of activated monocytes (being released together with IL-1), thereby indicating immune activation, it is also an endogenous inhibitor of pro-inflammatory IL-1 signaling [41]. Therefore, it is more appropriate to denote our results on cytokines/chemokines as indicating a generalized immune activation (M1, Th-1, Th-2 and Treg) in schizophrenia and MNP. As such, the positive association between IL-10 and eotaxin detected in our study may suggest that increased eotaxin levels accompanies the immune response in schizophrenia.

### 4.2. Immune activation and symptom dimensions of schizophrenia

The second major finding of this study is that the DAPS/PHEMN dimensions are highly associated with indicants of immune activation. Thus, the immune-defined cluster characterized by increased levels of cytokines/chemokines and TRYCATs explained around 42.8% of the variance in the overarchical “generalized psychopathology dimension” (consisting of all symptoms dimensions), while the zsIL1RA+MIP+IL10 index explained 19.2% and eotaxin 19.8% of the variance in this psychopathology dimension. Previously, it was detected that signs of immune activation including increased eotaxin levels are associated with specific symptom dimensions, e.g. positive symptoms [16]. Nevertheless, here we observed that all DAPS and PHEMN dimensions were highly significantly associated with the zsIL1RA+MIP+IL10 index, eotaxin and TRYCAT levels. Previously, we have discussed that activation of many different M1 and Th-1-associated pathways may translate into neurotoxic and cytotoxic effects [18,42,43], which could explain, at least in part, the association between immune activation and symptom dimensions. In addition, eotaxin may rapidly pass the blood-brain barrier to gain access to the brain and therefore increased serum eotaxin levels may be accompanied by eotaxin accumulation in CNS areas [44]. Neurons express both eotaxin and its receptor (CCR3) whereby the brain eotaxin-CCR3 system may exert physiological functions, including CCR3-associated neuroprotective and anti-inflammatory properties by promoting a Th2 phenotype [45,46]. Nevertheless, eotaxin has clear biphasic effects with physiological levels displaying neuroprotective activities and higher levels exerting neurotoxic effects. For example, eotaxin is associated with Alzheimer’s disease (AD) and age at onset of AD and additionally may affect hippocampal neurogenesis [44,47]. Therefore, our findings may suggest that increased eotaxin levels in schizophrenia play a role in symptom formation and consequently in shaping the MNP phenotype.

### 4.3. Association between immune activation and induction of the TRYCAT pathway

The third major finding of this study is that the zsIL1RA+MIP+IL10 index is significantly associated with TRYCAT pathway activation (as measured with the different TRYCAT ratios as well as with the total sum of all IgA responses to the 6 TRYCATs measured here). Previously, Schwieler et al. [48] found increased CSF IL-6, KA and kynurenine levels and suggested that increased IL-6 may be involved in IDO induction in schizophrenia. Our results indicate that, in schizophrenia, the TRYCAT pathway is activated by Th-1 and M1-related mechanisms via induction of IDO [11] and additionally that immune induced IDO activation is a hallmark of MNP. Importantly, immune activation (as measured with zIL1RA+MIP+IL10) was significantly associated with IgA levels directed against PA, XA and 3HK (the noxious TRYCATs) and not with AA or KA (the more protective TRYCATs). By inference, such data may indicate that immune activation may not only affect IDO, but also KMO (kynurenine monooxygenase), which activates production of 3HK, PA, XA and QA. It is indeed known that systemic immune activation induces a robust stimulation of brain KMO expression without any change in kynurenine aminotransferase II expression [49].

### 4.4. TRYCATs mediate the effects of immune activation on symptom dimensions

Here we established (using PLS analysis) that the effects of immune activation on the different symptom dimensions were completely mediated by increased activity of the TRYCAT pathway. It should be stressed that the effects of eotaxin on the symptom dimensions are not mediated via TRYCATs, resulting in complementary effects of increased TRYCAT and eotaxin levels in predicting schizophrenia phenomenology and thus the MNP phenotype. Previously, we have reviewed the many pathways by which neurotoxic TRYCATs may cause schizophrenia symptoms [1,6]. Thus, PA, XA and 3HK and QA have diverse cytotoxic, neurotoxic and excitotoxic effects and additionally cause inflammatory and oxidative responses which all together may induce adverse effects on neuronal functions. Of course, our results do not exclude the possibility that the antecedents of TRYCAT pathway activation (including increased M1, Th-1 and radical oxygen species activation) may contribute to the neurotoxic and cytotoxic effects of neuro-immune activation.

### 4.5. Effects of immune activation, including eotaxin and TRYCATs, on memory

The fourth major finding of this study is that the immune activation index and eotaxin have a strong impact on semantic and episodic memory and a more generalized deficit as well. Thus, 25.0% of the variance in cognitive CERAD tests was explained by these biomarkers, whereas both eotaxin and the immune activation index explained each around 18.0%. M1 and Th-1 cytokines have well known adverse effects on neurocognition, including IL-1β [30], IL-6 [50], IL-2 [51,52], IFN-γ [53] and IL-12 [54]. In addition, MIP-1 may impair hippocampal synaptic plasticity and thus may affect memory and learning [55]. MIP-1 binding to its receptor, CCR5 (chemokine receptor 5), causes accumulation of activated glial cells in the hippocampus [56]. Eotaxin is associated with reduced neurogenesis and impairments in hippocampal-mediated memory functions and learning [57]. Such mechanisms may explain the strong impact of eotaxin on episodic and semantic memory functions observed in schizophrenia patients.

Moreover, the current study detected that the effects of immune activation, but not eotaxin, on cognitive functions were completely mediated by TRYCATs. Phrased differently, increased severity of DAPS/PHEMN symptoms is related to the path from immune activation ➔ TRYCATs ➔ memory deficits ➔ symptom dimensions, and the path eotaxin ➔ memory deficits ➔ symptom dimensions. In this model, immune activation, increased levels of PA, XA and 3HK (and QA) together with eotaxin determine to a large extent memory impairments and the combined neuro-immune and neuro-cognitive disorders, in turn, determine to a large extent the different symptom dimensions and thus the MNP phenotype. Importantly, memory disorders, especially impairments in episodic memory, may at least in part explain some of the symptom dimensions of schizophrenia. Thus, “compromised cerebral functions”, preceding schizophrenia symptoms, may cause cognitive impairments [58,59], while abnormal learning processes and attentional impairments may generate psychosis and false memories [60]. False memory creation, another hallmark of MNP, may also increase risk for depressive symptoms [61].

### 4.6. New model of schizophrenia

Based on our findings we propose to revise previous theoretical cognitive models of schizophrenia [58,62,63]. Firstly, Harvey et al. [58] considered that negative symptoms and cognitive deficits are identical features or that both cognition and negative symptoms are distinct dimensions with a different pathophysiology. Nevertheless, our findings show that cognitive deficits and DAPS/PHEMN symptoms are highly interrelated phenomena that are largely determined by neuro-immune pathways. Secondly, Frith’s model considers that positive and negative symptoms are associated with impairments in self-awareness or consciousness [62], while Orellana and Slachdevsly [63] proposed that cognitive impairments (especially executive functions), which are the consequence of dysfunctions in brain neurocircuitry, underpin schizophrenia symptomatology. However, here we propose a revised theory, namely that activated immune pathways (mainly M1 and Th-1) induce IDO and KMO activity thereby increasing the production of noxious TRYCATs (PA, XA, 3HK and QA) and that these neuro-immune pathways coupled with elevated eotaxin levels (related to Th-2 activation) impact episodic and semantic memory, which all together lead to DAPS/PHEMN symptoms and thus the SNP and MNP phenotypes of schizophrenia.

## Acknowledgements

The study was supported by the Asahi Glass Foundation, Chulalongkorn University Centenary Academic Development Project and Ratchadapiseksompotch Fund, Faculty of Medicine, Chulalongkorn University, grant number RA60/042.

## Author’s contributions

All the contributing authors have participated in the manuscript. MM and BK designed the study. BK recruited patients and completed diagnostic interviews and rating scale measurements. MM carried out the statistical analyses. ST carried out the cognitive tests. SS performed all assays. All authors (BK, ST, SS, AC, AC and MM) contributed to interpretation of the data and writing of the manuscript. All authors approved the final version of the manuscript.

